# Interhemispheric callosal projections enforce response fidelity and frequency tuning in auditory cortex

**DOI:** 10.1101/2020.05.05.079012

**Authors:** Bernard J. Slater, Jeffry S. Isaacson

## Abstract

Sensory cortical areas receive glutamatergic callosal projections that link information processing between brain hemispheres. However, the role of interhemispheric projections in sensory processing is unclear. Here we use single unit recordings and optogenetic manipulations in awake mice to probe how callosal inputs modulate spontaneous and tone-evoked activity in primary auditory cortex (A1). Although activation of callosal fibers increased firing of some pyramidal cells, the majority of responsive cells were suppressed. In contrast, callosal stimulation consistently increased fast spiking (FS) cell activity and brain slice recordings indicated that parvalbumin (PV)-expressing cells receive stronger callosal input than pyramidal cells or other interneuron subtypes. *In vivo* silencing of the contralateral cortex revealed that callosal inputs linearly modulate tone-evoked pyramidal cell activity via both multiplicative and subtractive operations. These results suggest that callosal input regulates both the salience and tuning sharpness of tone responses in A1 via PV cell-mediated feedforward inhibition.

## Introduction

Cortical sensory representations driven by thalamic inputs are strongly influenced by local intracortical circuits and long range projections including interhemispheric inputs (Carrasco et al., 2013a, 2015; Cerri et al., 2010; Lee et al., 2019; Li et al., 2013; Lien and Scanziani, 2013; Schmidt et al., 2010; Wunderle et al., 2015). In most sensory systems there is an early decussation such that each hemifield of a sensory modality is primarily represented in the contralateral hemisphere of the brain. However, sensory areas for a particular modality in both cortices are linked to each other via interhemispheric projections from axons within the corpus callosum. These long range, cortico-cortical projections contact a majority of neurons in both supra- and infragranular layers (Carr and Sesack, 1998; Petreanu et al., 2007; Wise and Jones, 1976), but their postsynaptic targets and degree of connectivity vary for different sensory cortical areas (Harris et al., 2019). The differences in callosal connectivity with pyramidal cells and local interneurons is reflected in previous studies indicating that activation of callosal inputs can drive excitation and/or inhibition in cortical circuits (Anastasiades et al., 2018; Karayannis et al., 2007; Lee et al., 2014; Rock and Apicella, 2015). Although recent studies have begun to characterize the functional properties of interhemispheric cortical projections, how these pathways contribute to sensory coding *in vivo* is not well understood.

Unlike the visual and somatosensory cortices where interhemispheric inputs are relegated to hemifield overlap areas (Choudhury et al., 1965; Conti et al., 1986; Ebner and Myers, 1965; Hubel and Wiesel, 1967; Manzoni et al., 1989), callosal inputs are widespread across the tonotopically-organized primary auditory cortex (Code and Winer, 1985, 1986; Hackett and Phillips, 2011). Furthermore, anatomical studies in mammals indicate that callosal projections between primary auditory areas are “homotypic”: projections arising from a particular tonotopic region in one cortex map onto the corresponding frequency space within the contralateral cortex (Diamond et al., 1968; Imig and Brugge, 1978; Lee and Winer, 2008; Rouiller et al., 1991). Although callosal inputs arise from the axons of pyramidal cells in the opposite cortex, this pathway may not simply lead to cortical excitation. Indeed, in anesthetized ferrets, electrical stimulation of callosal inputs caused a variety of effects on sound-evoked firing rates including enhancement, suppression, or a mixture of the two (Kitzes and Doherty, 1994). Furthermore, intracellular recordings in A1 of anesthetized cats found that electrical stimulation in contralateral A1 elicited excitatory postsynaptic potentials that were often followed by inhibitory postsynaptic potentials (Mitani and Shimokouchi, 1985). These findings are consistent with a recent brain slice study indicating that A1 callosal inputs drive strong activation of layer 5 (L5) PV cells that mediate feedforward inhibition of pyramidal cells (Rock and Apicella, 2015). Despite these results suggesting a potential inhibitory influence of callosal inputs in auditory processing, removing interhemispheric input in anesthetized cats using cortical cooling was found to reduce sound-evoked activity in contralateral primary cortex (Carrasco et al., 2013a). However, anesthesia itself strongly influences spontaneous and sensory-evoked activity in sensory cortex (Harris and Thiele, 2011; Kato et al., 2015) and it is unclear how callosal input modulates A1 sensory processing in the awake state.

In this study, we use linear silicon probes spanning cortical layers to record spontaneous and tone-evoked single unit activity in A1 of awake, head-fixed mice. We express channelrhodopsin-2 (ChR2) in callosal fibers to study how their local activation modulates activity *in vivo* and identify the local circuits driven by callosal input in brain slice recordings. Finally, we use ChR2 in GABAergic interneurons to acutely suppress activity in one hemisphere while recording tone-evoked responses in contralateral A1 to show how the callosal pathway modulates cortical sensory processing.

## Results

We first studied how local activation of callosal projections modulates cortical excitability by targeting injection of adeno-associated virus (AAV) expressing ChR2 to A1 of the left hemisphere (Fig. 1A). Dense expression of ChR2 in fibers within the left medial geniculate body (MGB) confirmed that injections targeted primary auditory cortex (Fig. 1A_2_). We inserted linear silicon electrodes in A1 of the right hemisphere to monitor single unit activity in the awake state. Post-hoc analysis of probe recording sites revealed callosal ChR2-expressing fibers distributed across all layers of A1 (Fig. 1A_2_). Trough to peak time and full width at half maximum of spike waveforms (Fig. 1B) were used to classify single units as regular spiking (principal cells) or fast spiking (presumptive PV-expressing interneurons).

**Figure 1.**
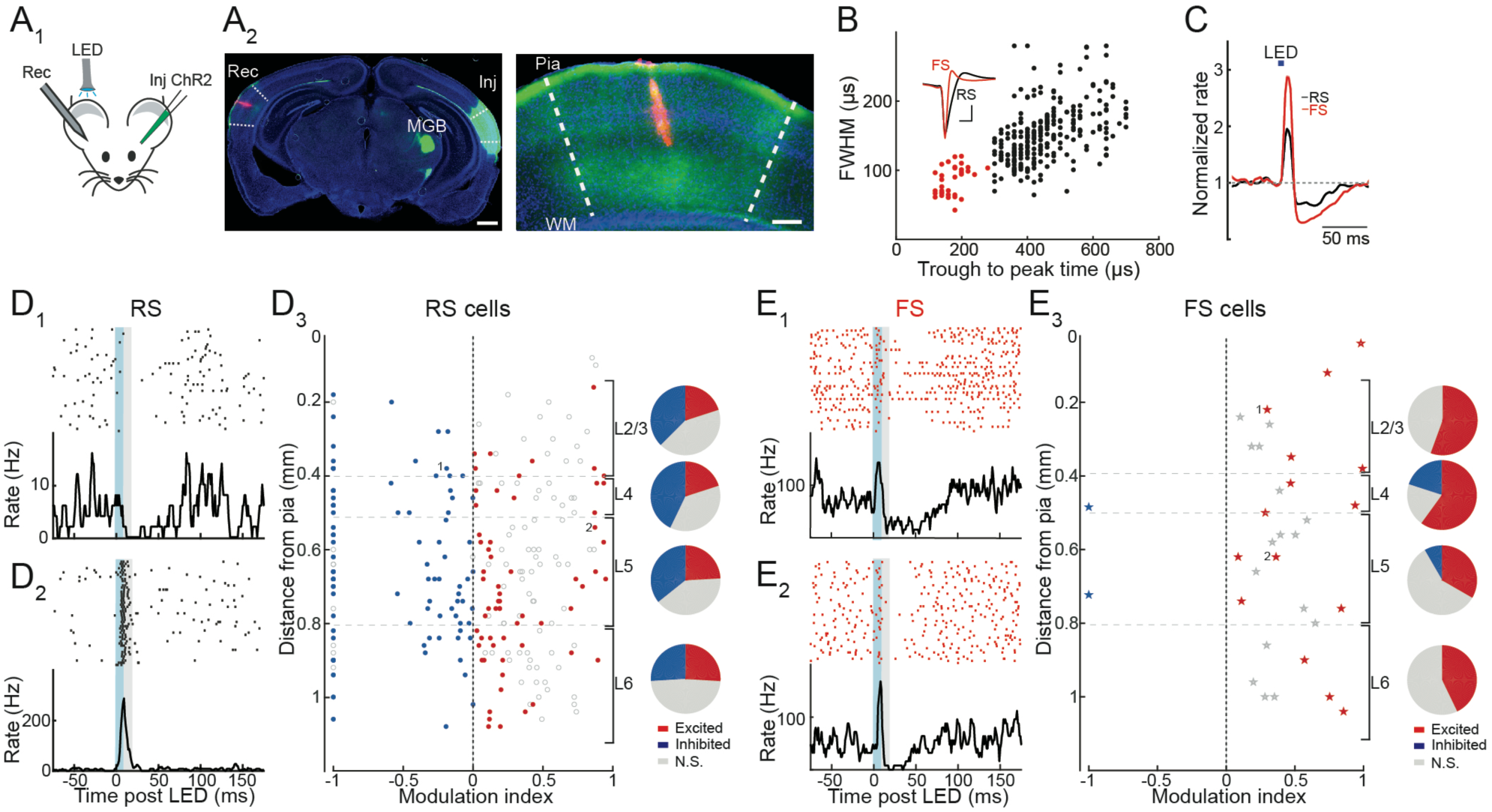
Optogenetic activation of cortical callosal inputs evokes excitation and inhibition in A1 of awake mice. A_1_, Experiment schematic. A_2_, Left, Coronal section showing ChR2 expression (green) within A1 of the injected left hemisphere (lnj) and DiI-labeled recording electrode tract (red) in contralateral A1 (Rec). Dense ChR2 expression is also present in the medial geniculate body (MGB) of the injected hemisphere. Scale bar = 1 mm. Right, Blow-up of recording site in the right hemisphere shows expression of ChR2-expressing fibers throughout all cortical layers. WM = white matter, scale bar = 250 µm. B, FS (red) and RS (black) units are identified by plotting spike trough to peak time vs. full width at half max (FWHM). Inset, average waveforms of FS and RS units. Scale bar = 250 µs, 20 µV. C, Average normalized peri-stimulus time histogram (PSTH) of RS (black) and FS (red) units shows that brief LED illumination (bar) drives a transient increase followed by a decrease in firing rate. D, Activation of callosal inputs increases activity of some RS cells, but inhibition is more widespread. D_1_, Individual RS unit spike raster and STH showing that ChR2 activation of callosal fibers (blue shading) inhibits firing. Grey shading indicates measurement period used to calculate modulation index. D_2_, RS unit strongly activated by callosal input. D_3_, Left, modulation index of units significantly activated (red) or inhibited (blue) across all layers. Open circles indicate units without significant effect and points marked 1 and 2 represent units in D_1_ and D_2_, respectively. Right, pie charts indicate proportion of units excited (red), inhibited (blue), or not significantly modulated (grey) in each layer. E, Activation of callosal inputs activates FS cells across all layers. Two representative FS units are plotted in E_1_ and E_2_. E_3_, modulation index of FS units across all cell layers are illustrated as for RS cells in D_3_.

We used brief (5 ms) LED illumination (470 nm) of the recording site to activate callosal inputs. On average, callosal stimulation caused a biphasic response in both RS (n = 264) and FS (n = 33, n = 7 mice) cells: a rapid increase in firing rate followed by a decrease in firing that returned to baseline over 50-100 ms (Fig. 1C). However, individual RS cells in the same experiments responded quite differently from each other: some cells were transiently excited by callosal stimulation, while others were exclusively inhibited (Fig. 1D_1,2_). We used a modulation index (Methods) to quantify early changes in firing (within 10 ms of callosal LED stimulation). We found that RS cells were more likely to be significantly inhibited than excited (Fig. 1D_3_, p < 0.05, sign test) in layers 2/3 (L2/3), 4 (L4) and 5, while cells were equally likely to be excited or inhibited in layer 6 (inhibited vs. excited, L2/3: 38 vs. 20% (n = 23 responding units), L4: 43 vs. 20% (n = 22), L5: 36 vs 24% (n=67), layer 6 (L6): 26% for each (n=40). In contrast, FS cells were much more likely to be significantly excited than inhibited by callosal stimulation across all layers (n = 15 excited vs. 2 inhibited, Fig. 1E). Together, these *in vivo* results indicate that while a subset of pyramidal cells are directly excited by callosal inputs, interhemispheric projections cause a widespread suppression of pyramidal cell activity. The rapid increase in FS cell firing evoked by activation of callosal inputs suggests that principal cell suppression arises from PV cell-mediated feedforward inhibition.

We next used voltage clamp recordings in brain slices to better understand the layer and cell type specificity of callosal input. We first examined the relative strength of callosal input onto PV and pyramidal cells. PV-Cre mice were crossed to a td-Tomato reporter line (Ai14) to target whole-cell recordings of visually identified PV cells and AAV-ChR2 was injected into A1 of the left cortex. We measured responses using simultaneously recorded pairs of PV and pyramidal cells (Pyr) from L2/3 of the cortex contralateral to the injection (Fig. 2A_1_). At −70 mV (near the reversal potential for GABAergic inhibition), brief LED illumination (470 nm, 2-4 ms) elicited excitatory postsynaptic currents (EPSCs) that were much larger in PV than pyramidal cells (peak EPSC amplitude PV=628±80 pA, Pyr=168±50 pA, n=6 pairs, p=0.003, paired t-test). Depolarization to +10 mV (near the reversal potential for glutamatergic excitation), revealed callosal input-evoked inhibitory postsynaptic currents (IPSCs) in both cell types. IPSCs always followed EPSCs with a brief delay in pyramidal and PV cells (average latency 2.13±0.51 ms, n = 8, and 1.81 ±0.2 ms, n = 10, respectively) indicating that inhibition was evoked indirectly by callosal input in a feedforward fashion (Isaacson and Scanziani, 2011). The ratio of excitation to inhibition (E/I ratio) was also markedly smaller in pyramidal than PV cells in L2/3 (0.11±0.01 and 0.33±0.06, respectively, n = 5 pairs, p = 0.01, paired t-test). Similarly, recordings in pairs of L5 pyramidal and PV cells revealed stronger callosal excitation of PV cells (Fig. 2A_2_, peak EPSC amplitude PV = 1105±324 pA, Pyr = 197±60 pA, n=6 pairs, p=0.03, paired t-test), a smaller pyramidal cell E/I ratio (ratio PV = 0.46±0.08, Pyr = 0.11±0.02, n = 5 pairs, p = 0.007, paired t-test), and disynaptic IPSC latency (PV = 1.48 ±0.07 ms, n = 10, Pyr = 1.08±0.11 ms, n = 5). Interestingly, paired recordings of L2/3 and L5 PV cells indicated that PV cells in deeper cortical layers receive more callosal excitation (Fig. 2A_3_, peak EPSC amplitude L2/3 = 0.81±0.21 nA, L5 = 2.13±0.43 nA, n = 7 pairs, p = 0.03, paired t-test) and had a higher E/I ratio (L2/3 = 0.29±0.05, L5 = 0.54±0.07, n = 7 pairs, p = 0.03, paired t-test). These findings indicate that callosal projections drive stronger excitation of PV cells than pyramidal cells in both infra- and supragranular layers. Furthermore, activation of callosal input drives strong feedforward inhibition of principal cells in A1.

**Figure 2.**
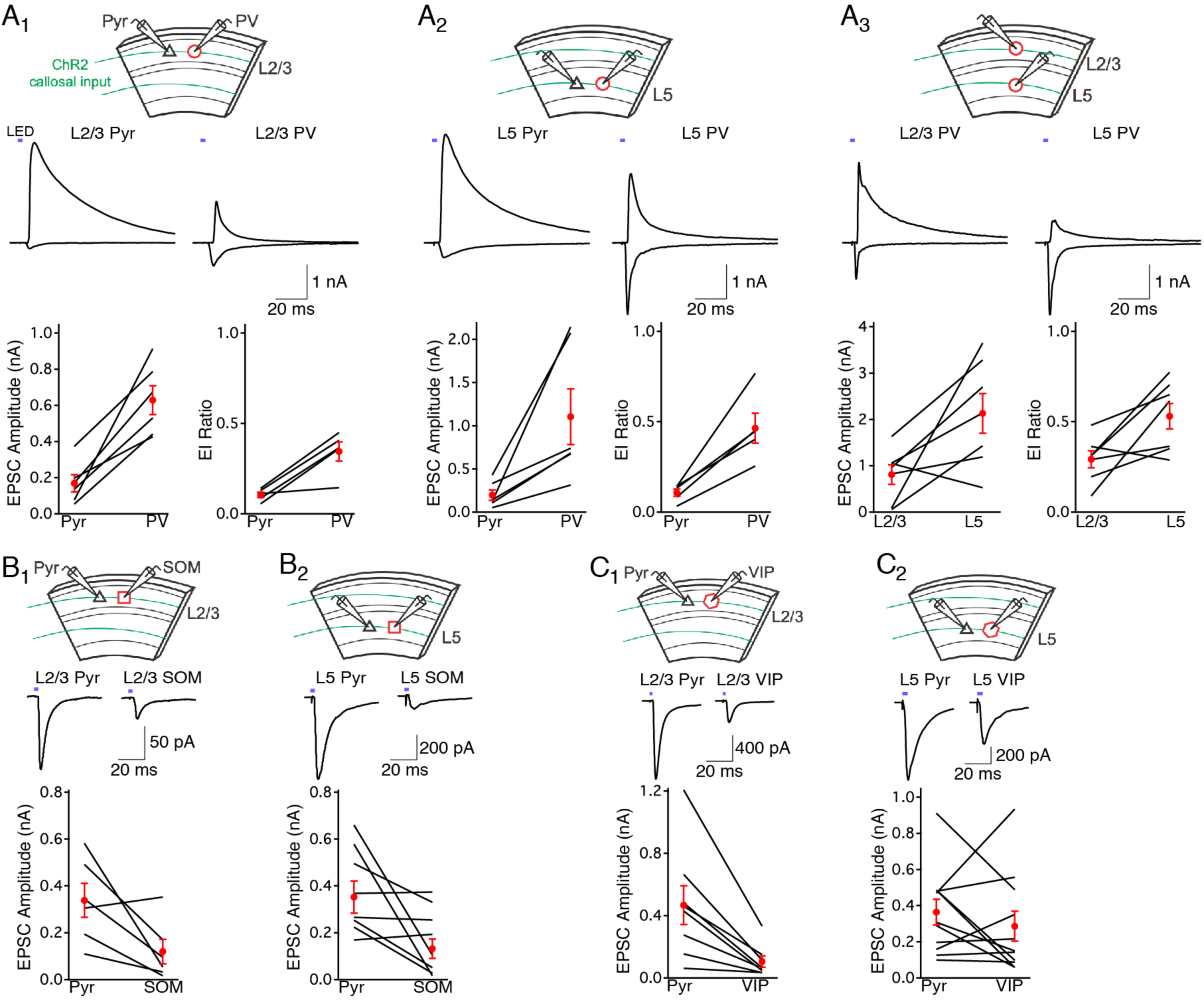
Cortical callosal inputs preferentially excite PV cells and drive strong feedforward inhibition. A_1_, L2/3 PV cells receive stronger callosal fiber-evoked EPSCs and have a larger E/I ratio than L2/3 pyramidal cells. Top, recording configuration. Middle, simultaneous voltage clamp recording of L2/3 pyramidal cell (Pyr) and PV cell showing EPSCs (inward currents, −70 mV) and IPSCs (outward currents, +10 mV) evoked by brief LED illumination (blue bars) of ChR2-expressing callosal fibers. Bottom, summary of EPSC peak amplitudes and E/I ratios for recorded pairs. Black lines, individual cell pairs. Red circles, mean ±SEM. A_2_, L5 PV cells receive stronger callosal fiber-evoked EPSCs and have a larger E/I ratio than L5 pyramidal cells. A_3_, L5 PV cells receive stronger callosal fiber-evoked EPSCs and have a larger E/I ratio than L2/3 PV cells. B, SOM cells in L2/3 (B_1_) and L5 (B_2_) receive weaker callosal fiber-evoked EPSCs than neighboring pyramidal cells. C, VIP cells in L2/3 (C_1_) receive weaker callosal fiber-evoked EPSCs than neighboring pyramidal cells. The strength of callosal input-evoked EPSCs in L5 VIP cells (C_2_) and pyramidal cells are similar.

Are PV cells unique or do all classes of interneurons receive stronger callosal input than pyramidal cells? To address this, we recorded callosal input-evoked EPSCs onto pairs of pyramidal cells and td-Tomato labeled somatostatin (SOM)- or vasoactive intestinal polypeptide (VIP)-expressing interneurons using SOM- and VIP-Cre mice. Activation of ChR2-expressing callosal inputs evoked EPSCs that were markedly weaker in SOM cells compared to pyramidal cells in both L2/3 (Fig. 2B_1_, peak EPSC amplitude SOM = 120±52 pA, Pyr = 338±73 pA, n = 6 pairs, p = 0.04, paired t-test) and L5 (Fig. 2B_2_, SOM = 132±41 pA, Pyr = 352±69 pA, n = 8 pairs, p = 0.03, paired t-test). Callosal EPSCs were much weaker in VIP cells compared to pyramidal cells in L2/3 (Fig. 2C_1_, peak EPSC amplitude VIP = 105±36 pA, Pyr = 467±126 pA, n = 8 pairs, p = 0.006, paired t-test) while responses were roughly similar in L5 (Fig. 2C_2_, VIP = 285±83 pA, Pyr = 364±71 pA, n = 11 pairs, p = 0.37, paired t-test). The relatively weak callosal-evoked EPSCs in SOM and VIP interneurons suggest that they are not a major target of interhemispheric input.

To directly examine the functional role of interhemispheric input *in vivo*, we recorded from A1 in awake mice while optogenetically suppressing activity in the contralateral auditory cortex. We injected AAV-FLEX-ChR2 (Atasoy et al., 2008) in the left cortex of Gad2-Cre mice to express ChR2 in GABAergic interneurons (Fig. 3A_1_, Kato et al., 2015). Recordings in the injected cortex confirmed that LED illumination (20 Hz train of 10 ms pulses) drove firing of FS cells (Fig. 3A_2_) while RS cell activity was largely abolished (Fig. 3A_3_,_4_). We next monitored spontaneous activity in A1 of the right hemisphere while silencing contralateral A1 (Fig. 3B_1_). Although it has been suggested that GABAergic interneurons in auditory cortex can make interhemispheric projections (Rock et al., 2018), we did not observe ChR2-expressing fibers in A1 contralateral to the AAV-injected cortex (Fig. 3B_2_). On average, silencing A1 in the left hemisphere caused a transient decrease in firing followed by an increase in activity in RS and FS cells in contralateral A1 (Fig. 3B_3_, n = 494 RS, 76 FS, n = 19 mice). However, individual cells responded differently to contralateral silencing depending on cortical layer. LED-responsive RS cells in layers 2/3, 4 and 5 primarily increased their firing during cortical silencing (excited vs. inhibited: 21 vs. 6%, n = 90 responding units), while L6 RS cells were typically inhibited (excited vs. inhibited: 8 vs. 29%, n = 23 responding units). Similarly, FS cells in layers 2/3 and 4 were primarily excited during cortical silencing (excited vs. inhibited: 49 vs. 24%, n = 31 responding units) while those in layers 5 and 6 were more likely to be suppressed (excited vs. inhibited: 11 vs. 43%, n = 20 responding units). These results indicate that spontaneous firing in L6 RS cells and deep layer FS cells is dependent on callosal input. The increase in firing in upper layers during cortical silencing is likely to reflect network effects associated with the withdrawal of deep layer RS and FS cell activity.

**Figure 3.**
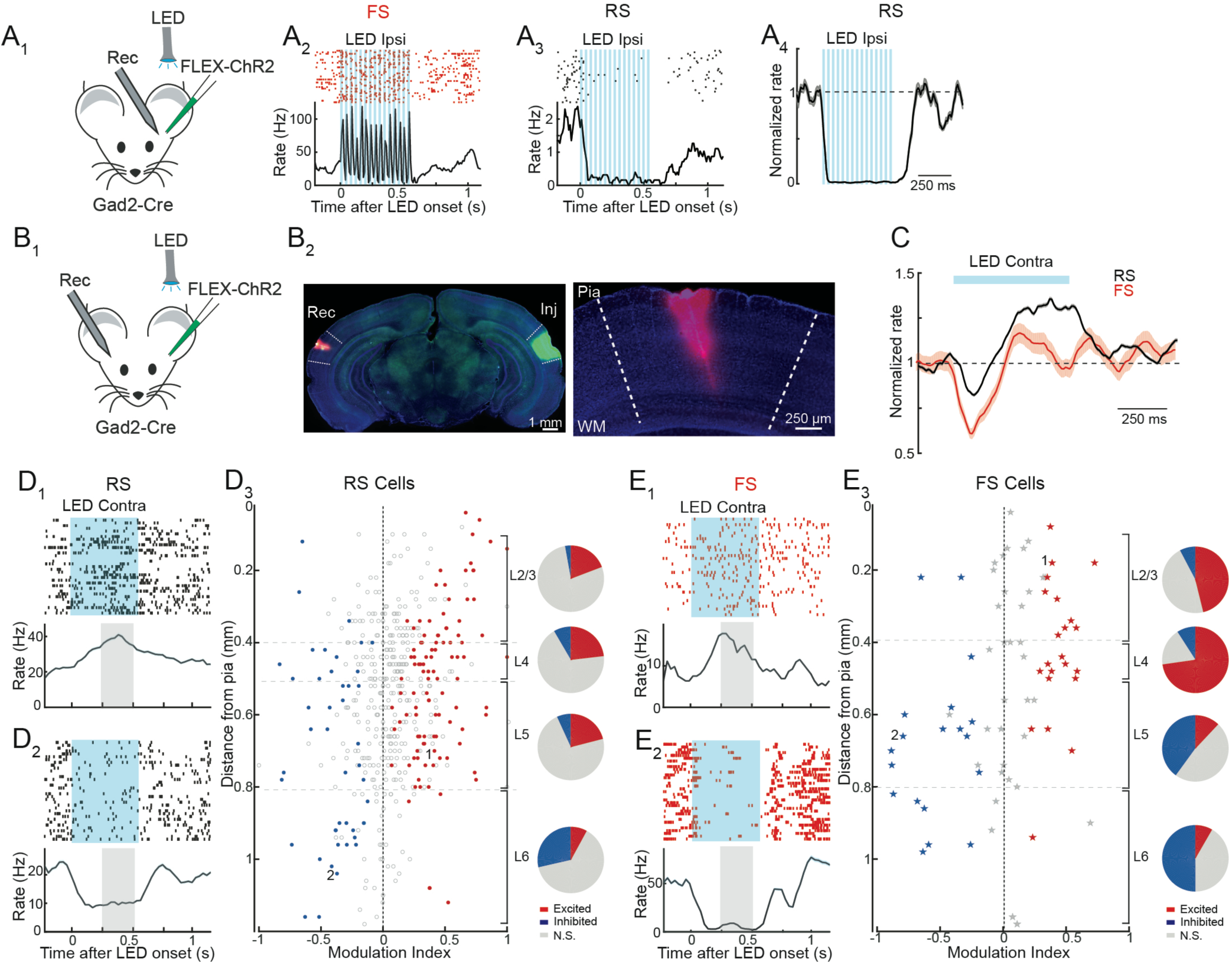
Acute optogenetic silencing of interhemispheric cortical input causes a sustained increase in spontaneous activity in most layers of A1. A, Local activation of ChR2-expressing interneurons silences RS cell activity. A_1_, recording configuration. A_2_, spike raster (top) and PSTH (bottom) show strong activation of a representative FS unit by an ipsilateral LED pulse train (blue bars). A_3_, spike raster (top) and PSTH (bottom) show strong suppression of simultaneously recorded RS unit. A_4_, summary of ipsilateral LED-evoked suppression of RS activity (n = 34 units, 2 mice). B, Activation of ChR2-expressing interneurons in one hemisphere leads to transient inhibition followed by excitation in contralateral A1. B_1_, recording configuration. B_2_, Left, Coronal section showing ChR2 expression (green) within A1 of the injected left hemisphere (lnj) and DiI-labeled recording electrode tract (red) in contralateral A1 (Rec). Right, Blow-up of recording site. WM = white matter. C, Average normalized PSTH of RS (black) and FS (red) units shows that sustained LED illumination (bar) drives transient decrease and sustained increase in firing. Shading, ±SEM. D, Inactivation of A1 causes sustained increase in activity of RS units in layers 1-5 of contralateral A1. D_1_, individual L5 RS unit spike raster and PSTH showing that silencing contralateral A1 (blue shading) enhances firing. Grey shading indicates measurement period used to calculate modulation index. D_2_, L6 RS unit with sustained suppression during silencing of contralateral A1. D_3_, Left, modulation index of units significantly activated (red) or inhibited (blue) across all layers. Open circles indicate units without significant effect and cells marked 1 and 2 represent units in D_1_ and D_2_, respectively. Right, pie charts indicate proportion of units excited (red), inhibited (blue), or not significantly modulated (grey) in each layer. E, Silencing contralateral A1 causes a rapid and sustained decrease in firing in deep layer FS cells, as well as a sustained firing increase in upper layer FS cells. Representative L2/3 and L5 FS unit are plotted in E_1_ and E_2_, respectively. E_3_, modulation index of FS units across all cell layers are illustrated as in D_3_.

We next examined how silencing contralateral cortex modulates tone-evoked activity of RS cells in A1. The right ear was occluded and pure tones (9 log-spaced frequencies, 4-60 kHz, 250 ms, 60 dB) were delivered to the left ear during optogenetic silencing of the left hemisphere on interleaved trials (Fig. 4A, tone onset 250 ms following start of LED illumination). RS cells recorded from right A1 were frequency-tuned (Fig. 4B) such that particular frequencies drove strong firing (“preferred tones”) while others evoked weak responses (“non-preferred tones”). Interestingly, the effects of cortical silencing on RS cell activity were dependent on the strength of tone-evoked responses. Firing rates during non-preferred tones were enhanced by contralateral silencing, while firing evoked by preferred tones were largely unaffected or reduced (Fig. 4B_1_, 4D_2_). This effect could be described by a simple linear transformation: firing rates during tones with vs. without LED-induced silencing could be fit by a line with a slope < 1 and y-intercept > 0 (Fig. 4B_2_). In other words, removing callosal input had both an additive and divisive action on A1 tone responses. The effects of contralateral cortical silencing were uniformly divisive across all cortical layers (Fig. 4C_1_) while additive effects were prominent in all but L6 (Fig 4C_2_). Together, these results suggest that callosal input normally regulates sound-evoked responses via multiplicative and subtractive effects.

**Figure 4.**
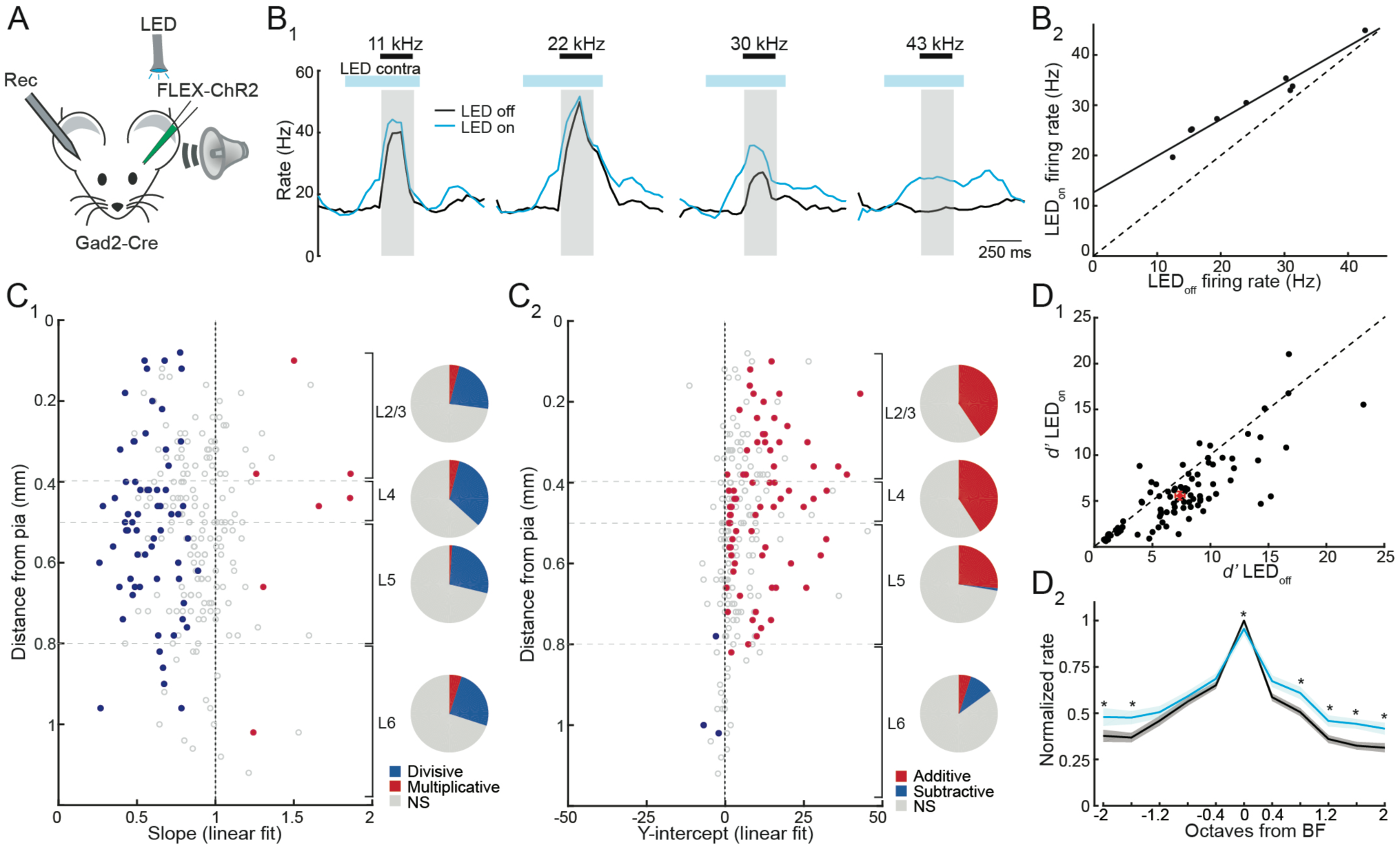
Silencing interhemispheric cortical input degrades the fidelity and frequency tuning of tone-evoked responses in A1. A, recording configuration. B, Silencing contralateral A1 linearly modulates tone evoked activity via a combination of additive and divisive operations. B_1_, PSTHs of tone-evoked responses from a representative RS unit to four frequencies (black bars) under control conditions (black line) and during contralateral silencing (blue line) on interleaved trials. Blue bars, LED pulse train. Grey, measurement windows for tone-evoked firing rate. B_2_, plot of firing rates during tones (n = 9 frequencies) with the LED on vs. LED off of the cell in B. Line is linear fit: slope= 0.73, y-intercept = 12.63, r^2^ = 0.96. C, Silencing callosal input exerts divisive and additive actions on tone-evoked activity across cortical layers. C_1_, Slopes derived from linear fits to individual RS units with significant tone-evoked activity in each cortical layer. Blue circles, slope significantly <1. Red circles, slope significantly >1. Open circles, no significant change in slope. Pie charts represent fraction of cells in each layer with divisive (blue, slope <1), multiplicative (red, slope>1), or no significant effect (grey, NS). C_2_, Y-intercepts derived from linear fits to same RS units in C_1_. Blue circles, y-intercept significantly <0. Red circles, y-intercept significantly >0. Open circles, y-intercept not significantly different from 0. Pie charts represent fraction of cells in each layer with additive (red, y-intercept >0), subtractive (blue, y-intercept <0), or no significant effect (grey, NS). D_1_, d’ of RS units with LED off vs. LED on shows that cortical silencing reduces response detectability. D_2_, Cortical silencing “flattens” frequency tuning curves. Average tuning curves of RS units centered to their BF under control conditions (black) and during contralateral cortical silencing (blue). Asterisks indicate frequencies with significant difference (paired t-test, Holm-Bonferroni corrected).

Divisive/multiplicative operations exert gain control of neural responses while subtractive/additive operations modulate response fidelity via changes in variability associated with stimulus-independent (“background”) activity (Isaacson and Scanziani, 2011; Silver, 2010). Both the increase in spontaneous activity and additive effects on tone responses during contralateral cortical silencing suggest that callosal inputs enforce response fidelity. To address this possibility, we computed the discriminability index (*d’*, Methods), a measure of response reliability from signal detection theory (Duguid et al., 2012; Sturgill and Isaacson, 2015; Tolhurst et al., 1983) with and without contralateral cortical silencing. Optogenetic cortical inactivation significantly reduced the discriminability of tone-evoked activity (Fig. 4D_1_ *d′*_LED-off_ = 7.37 ± 0.45, *d′*_LED-on_ = 5.58 ± 0.41, *n* = 124, *P* < 0.001, t-test) indicating that callosal input normally serves to enhance the representation of tone responses relative to spontaneous activity in A1.

We examined how callosal input modulates the shape of frequency tuning curves by normalizing cell responses to their best frequency (BF, tone eliciting strongest increase in firing) under control conditions. Silencing contralateral cortex caused a small decrease in the amplitude of responses at BF (Fig. 4D_2_, p = 0.01, t-test), consistent with the divisive effect we observed on input-output relationships (Fig. 4C). However, due to its additive action, cortical silencing also increased responses to non-preferred frequencies. The net effect is thus a “flattening” of the population frequency tuning curve (Fig. 4D_2_). Thus, in addition to regulating response fidelity, callosal inputs normally play an important role in enforcing the sharpness of frequency tuning in A1.

## Discussion

We show that activating interhemispheric callosal projections can inhibit pyramidal cells in all layers of A1 in awake mice. These findings are consistent with slice recordings indicating that callosal inputs evoke strong feedforward inhibition of pyramidal cells in supra- and infragranular layers. This feedforward inhibition likely reflects the recruitment of PV cells, which receive stronger callosal excitation than SOM or VIP cells in upper and lower cortical layers. In loss-of-function experiments, acute *in vivo* silencing of contralateral cortex increased pyramidal cell spontaneous activity in all but L6. Finally, we used tone-evoked activity to show that cortical silencing linearly transforms A1 input-output relationships via subtractive and divisive operations. This indicates that interhemispheric projections normally enhance the salience of tone representations (by regulating signal to noise ratio) and sharpen frequency tuning in primary auditory cortex.

It is well established that callosal inputs make direct excitatory connections onto cortical pyramidal cells (Anastasiades et al., 2018; Karayannis et al., 2007; Lee et al., 2014, 2019; Petreanu et al., 2007; Rock and Apicella, 2015) and drive disynaptic feedforward inhibition via contacts onto local GABAergic interneurons (Anastasiades et al., 2018; Karayannis et al., 2007; Rock and Apicella, 2015). Indeed, we found that brief activation of callosal fibers drives a biphasic increase and decrease in the firing of RS and FS cells in awake mice. Surprisingly, individual RS cells across all cortical layers were more likely to be inhibited than excited by callosal stimulation. In contrast, FS cells were more routinely activated, suggesting that the suppressive effects of callosal stimulation on RS cell firing are due to widespread PV cell-mediated feedforward inhibition. Consistent with this idea, brain slice recordings revealed that PV cells receive more callosal input than neighboring pyramidal cells or other interneuron subtypes and deep layer PV cells received ~2X stronger input than L2/3 PV cells.

Previous studies in sensory cortical areas have used callosal sectioning (Engel et al., 1991; Payne et al., 1980) or reversible cortical cooling to probe the functional role of callosal inputs in anesthetized animals (Carrasco et al., 2013, 2015; Cerri et al., 2010; Schmidt et al., 2010; Wunderle et al., 2015). We show in awake mice that acute optogenetic silencing has heterogeneous effects on spontaneous activity: although a subset of RS cells shows a rapid and sustained decrease in activity, the majority of cells responded with a slow sustained increase in firing. The simplest interpretation of these results is that decreases in activity reflect the withdrawal of direct excitatory callosal input onto particular cells, while paradoxical increases in firing reflect indirect network effects. Increases in firing are most likely due to a reduction in inhibition provided by PV cells. Indeed, we observed that the spontaneous firing of deep layer PV cells was strongly suppressed during contralateral cortical silencing. This suggests that much of the tonic activity of deep layer PV cells is driven by interhemispheric input. Deep layer interneurons have recently been shown to project axons through all cortical layers towards the pia (Bortone et al., 2014; Frandolig et al., 2019). It is possible that interlaminar projections from deep layer PV interneurons mediate the indirect network effects underlying principal cell excitation following withdrawal of callosal input.

In contrast to previous work in auditory cortex of anesthetized animals (Carrasco et al., 2013b, 2015), we did not observe a simple reduction in the strength of tone-evoked responses during contralateral silencing in the awake state. Rather, input-output plots of tone-evoked firing were linearly transformed in a divisive and additive fashion, reflecting both the withdrawal of direct callosal excitatory input on pyramidal cells and reduction in feedforward inhibition. Higher spontaneous activity and stronger inhibition in the awake state are likely to underlie these differences (Haider et al., 2013; Kato et al., 2015). In terms of frequency tuning, this led to a small reduction in responses to BF while responses to flanking non-preferred frequencies were enhanced. Thus, in addition to enhancing the discriminability of sound-evoked responses by maintaining a high signal to noise ratio, callosal inputs sharpen frequency tuning in primary auditory cortex. These findings are in agreement with previous studies indicating that interhemispheric connections modulate the specificity of sensory-evoked activity in visual (Hubel and Wiesel, 1967; Schmidt et al., 2010; Wunderle et al., 2015) and somatosensory cortex (Clarey et al., 1996). In future, it will be useful to determine how callosal input contributes to binaural cortical sound representations and auditory-directed behaviors such as sound localization.

## Methods

Mice (8–16 weeks old for *in vivo* recordings, 3-5 weeks old for *in vitro* recordings, Emx1-Cre (Jackson Laboratories No. 05638), Gad2-Cre (Jackson Laboratories No. 019022), PV-cre (Jackson Laboratories No. 017320), SOM-Cre (Jackson Laboratories No. 010708), VIP-cre (Jackson Laboratories No. 010908), tdTomato reporter (Ai14, Jackson Laboratories No. 00914), and wild-type C57Bl6 mice were housed with a 12:12 h reversed light cycle. All *in vivo* experiments were performed during the dark period. All procedures were in accordance with protocols approved by the University of California San Diego Institutional Animal Care and Use Committee and guidelines of the National Institutes of Health.

### Surgical preparation

For *in vivo* electrophysiology experiments, 2-3 weeks prior to head-bar implantation and habituation to head fixation, mice were anesthetized with isoflurane, and the skull above left A1 identified by intrinsic imaging (Kato et al., 2015). Viruses (AAV9-hSyn-hChR2(H134R)-eYFP-WPRE-hGH or AAV9-Ef1α-DIO-hChR2(h134R)-YFP-WPRE-hGHpA, UPenn) were injected (50 nL) using beveled pipettes (Nanoject II, Drummond) at three sites spanning A1 at depths of 0.25–0.75 mm. After injections, mice received dexamethasone (2 mg/kg), buprenorphine (0.1 mg/kg) and baytril (10 mg/kg) prior to returning to their home cage. 2-3 days prior to *in vivo* recording, a head bar was implanted and right A1 was identified using intrinsic imaging.

For *in vitro* recordings, neonatal mice (postnatal day 0–2) were anesthetized and virus injection sites targeting the auditory cortex, which were determined based on landmarks including the superficial temporal vein. AAV9-hSyn-hChR2(H134R)-eYFP-WPRE-hGH (ChR2-YFP) was injected (23 nL) using beveled pipettes (Nanoject II, Drummond) at three injection sites at depths of 0.2–0.6 mm. Experiments were performed >2 weeks after injection.

### Extracellular recordings

A 32-(Neuronexus) or 64-(Cambridge Neurotech) channel silicon probe was used for extracellular recordings. Signals were recorded using an Intan RHD2000 and digitized at 20 kHz using Open Ephys (Siegle et al., 2017). Spikes were sorted using Kilosort (Pachitariu et al., 2016), followed by manual curation in phy (Rossant et al., 2016) to obtain single units used for analyses. Cells were excluded from analysis if they did not maintain consistent firing and amplitude throughout recording, and a firing rate of at least 1Hz. The probe was coated in DiI to verify probe track for depth of recording as well as recording location. Current source density (Pettersen et al., 2006) coupled with anatomical verification of probe track was used to identify laminar single unit locations. For all recordings spike waveforms were obtained from the lead with the largest amplitude template, these were then averaged to obtain an average spike waveform. Units were classified as fast spiking if their average spike waveform had a trough to peak time of less than 300 µs and a full-width at half max of less than 125 µs.

A fiber-coupled LED (470 nm, 20 mW, 0.4 mm fiber, 0.48 N.A., Thorlabs) was positioned within 1-2 mm of the exposed cortical surface for activating ChR2-expressing callosal fibers or ipsilateral cortical silencing. For experiments using contralateral silencing, the skull over right auditory cortex was exposed and covered with cyanoacrylate glue before the LED fiber was positioned at the skull surface. Callosal fiber activation was achieved using a single 5 ms flash. For cortical silencing in Gad2-cre mice expressing ChR2 we used a train of 10 ms light pulses (510 ms, 20 Hz) to activate inhibitory interneurons.

Immediately prior to recording, mice were anesthetized and the ear canal contralateral to the recording was filled with cyanoacrylate glue to occlude the ear. To prepare the recording site, a well filled with artificial cerebrospinal fluid (aCSF, in millimoles: 142 NaCl, 5 KCl, 10 glucose, 10 HEPES, 3.1 CaCl_2_, 1.3 MgCl_2_, pH 7.4, 310 mOsm) was constructed around the recording site and a small (<0.3 mm) craniotomy was performed through thinned skull. Mice recovered for >1hr before the start of recording. Pure tones logarithmically spaced between 4 kHz and 60 kHz were delivered via a calibrated free-field speaker (ES1, TDT) directed to the left ear. Tones were generated by software (BControl; http://brodylab.org) running on MATLAB (MathWorks) communicating with a real-time system (RTLinux). Tone frequencies (250 ms duration) were presented in a pseudo-random fashion and LED illumination was delivered on interleaved trials.

### In vitro electrophysiology

Patch-clamp recordings were performed using an upright microscope, 40X objective, and DIC optics. Recordings were made using a Multiclamp 700A amplifier (Molecular Devices), digitized at 20 kHz, and acquired and analyzed using AxographX software. For voltage-clamp recordings, pipettes (3–5 MΩ) contained (in mM): 130 D-gluconic acid, 130 CsOH, 5 NaCl, 10 HEPES, 10 EGTA, 12 phosphocreatine, 3 Mg-ATP, and 0.2 Na-GTP (pH 7.3). Series resistance was routinely <20 MΩ and continuously monitored. LED illumination (470 nm, Thorlabs) was delivered through the microscope objective.

### Analysis of in vivo data

For presentation of pooled neuronal responses, firing rates were normalized to the average baseline firing rate of each neuron 250 ms before the LED period. The analysis window for callosal terminal excitation was 10 ms from LED onset to capture both the initial excitation and recurrent inhibition. In contralateral A1 silencing experiments, the window for analysis was a 250 ms time period that started 250 ms after LED onset. All statistical tests were two sided and used a significance level of 0.05 (corrected for multiple comparisons where noted). Units were considered significantly modulated by the LED if the mean firing rate during the analysis window was different than that of the baseline period as determined by a Wilcoxon sign-rank test α = 0.05. Modulation index was calculated as [(mean firing rate in analysis window) – (mean firing rate during baseline period)]/[(mean firing rate in analysis window) + (mean firing rate during baseline period)]. Average modulation of units was tested for significance using a one sample t-test.

Sound responses were determined as significant at a given frequency if p<0.05 for a Wilcoxon rank sum test of firing rate over 250ms starting 10 ms after sound onset as compared to the same time period during interleaved trials with no tones (blank trials). A Holm-Bonferroni correction was used for multiple comparisons. Units were considered sound responsive if they responded to at least one tone frequency. Unit responses to a given frequency were averaged and these average responses were fit with a linear polynomial. RS units were included in analysis if they were sound responsive and had a linear fit with r^2^ > 0.25. Slope significance was determined using a 95% confidence interval for the linear fit, slopes were considered significantly modulated either divisively or multiplicatively if the upper bound was <1 or the lower bound was >1 respectively. Intercept significance was determined using a 95% confidence interval for the linear fit, intercepts were considered significantly modulated in either an additive or subtractive fashion where lower bound was >0 or the upper bound was <0 respectively. The discriminability index, *d′*, was calculated for the average of every LED modulated tone response as (mean Spikes_sound_ − mean Spikes_spontaneous_)/√ [0.5 × (σ^2^_sound_ + σ^2^_spontaneous_)]. Tone responses for a given unit were excluded if their tone response versus spontaneous firing rate *z*-score was <2. The *d′* values are presented as the mean of *d′* values for a given unit. To generate a frequency tuning curves, individual unit responses were average at each frequency. The responses were then centered to the best frequency (BF) chosen as the frequency which had the strongest tone response in the control condition for each unit. Significant modulation at each frequency by cortical inactivation was determined using a paired t-test followed by a Holm-Bonferroni correction for multiple comparisons.

## Acknowledgements

We are grateful to Chris Song, Bella Nguyen, and Elena Westeinde for technical support. This work was supported by NIH R01DC04682 and R01DC015239 to J.S.I. and F32DC017906 to B.J.S.

## References

Anastasiades, P.G., Marlin, J.J., and Carter, A.G. (2018). Cell-Type Specificity of Callosally Evoked Excitation and Feedforward Inhibition in the Prefrontal Cortex. Cell Rep. 22, 679–692.

Atasoy, D., Aponte, Y., Su, H.H., and Sternson, S.M. (2008). A FLEX switch targets Channelrhodopsin-2 to multiple cell types for imaging and long-range circuit mapping. J Neurosci 28, 7025–7030.

Bortone, D.S., Olsen, S.R., and Scanziani, M. (2014). Translaminar inhibitory cells recruited by layer 6 corticothalamic neurons suppress visual cortex. Neuron 82, 474–485.

Carr, D.B., and Sesack, S.R. (1998). Callosal terminals in the rat prefrontal cortex: Synaptic targets and association with GABA-immunoreactive structures. Synapse 29, 193–205.

Carrasco, A., Brown, T.A., Kok, M.A., Chabot, N., Kral, A., and Lomber, S.G. (2013a). Influence of Core Auditory Cortical Areas on Acoustically Evoked Activity in Contralateral Primary Auditory Cortex. J. Neurosci. 33, 776–789.

Carrasco, A., Brown, T.A., Kok, M.A., Chabot, N., Kral, A., and Lomber, S.G. (2013b). Influence of core auditory cortical areas on acoustically evoked activity in contralateral primary auditory cortex. J. Neurosci. 33, 776–789.

Carrasco, A., Kok, M.A., and Lomber, S.G. (2015). Effects of Core Auditory Cortex Deactivation on Neuronal Response to Simple and Complex Acoustic Signals in the Contralateral Anterior Auditory Field. Cereb. Cortex 25, 84–96.

Cerri, C., Restani, L., and Caleo, M. (2010). Callosal contribution to ocular dominance in rat primary visual cortex. Eur. J. Neurosci. 32, 1163–1169.

Choudhury, B.P., Whitteridge, D., and Wilson, M.E. (1965). THE FUNCTION OF THE CALLOSAL CONNECTIONS OF THE VISUAL CORTEX. Q. J. Exp. Physiol. Cogn. Med. Sci. 50, 214–219.

Clarey, J.C., Tweedale, R., and Calford, M.B. (1996). Interhemispheric Modulation of Somatosensory Receptive Fields: Evidence for Plasticity in Primary Somatosensory Cortex. Cereb. Cortex 6, 196–206.

Code, R.A., and Winer, J.A. (1985). Commissural neurons in layer III of cat primary auditory cortex (AI): Pyramidal and non-pyramidal cell input. J. Comp. Neurol. 242, 485–510.

Code, R.A., and Winer, J.A. (1986). Columnar organization and reciprocity of commissural connections in cat primary auditory cortex (AI). Hear. Res. 23, 205–222.

Conti, F., Fabri, M., and Manzoni, T. (1986). Bilateral Receptive Fields and Callosal Connectivity of the Body Midline Representation in the First Somatosensory Area of Primates. Somatosens. Res. 3, 273–289.

Diamond, I.T., Jones, E.G., and Powell, T.P.S. (1968). Interhemispheric fiber connections of the auditory cortex of the cat. Brain Res. 11, 177–193.

Duguid, I., Branco, T., London, M., Chadderton, P., and Häusser, M. (2012). Tonic inhibition enhances fidelity of sensory information transmission in the cerebellar cortex. J. Neurosci. 32, 11132–11143.

Ebner, F.F., and Myers, R.E. (1965). Distribution of corpus callosum and anterior commissure in cat and raccoon. J. Comp. Neurol. 124, 353–365.

Engel, A., Konig, P., Kreiter, A., and Singer, W. (1991). Interhemispheric synchronization of oscillatory neuronal responses in cat visual cortex. Science (80-.). 252, 1177–1179.

Frandolig, J.E., Matney, C.J., Lee, K., Kim, J., Chevée, M., Kim, S.-J., Bickert, A.A., and Brown, S.P. (2019). The Synaptic Organization of Layer 6 Circuits Reveals Inhibition as a Major Output of a Neocortical Sublamina. Cell Rep. 28, 3131–3143.e5.

Hackett, T.A., and Phillips, D.P. (2011). The commissural auditory system. In The Auditory Cortex, (Springer US), pp. 117–131.

Haider, B., Häusser, M., and Carandini, M. (2013). Inhibition dominates sensory responses in the awake cortex. Nature 493, 97–100.

Harris, K.D., and Thiele, A. (2011). Cortical state and attention. Nat. Rev. Neurosci. 12, 509–523.

Harris, J.A., Mihalas, S., Hirokawa, K.E., Whitesell, J.D., Choi, H., Bernard, A., Bohn, P., Caldejon, S., Casal, L., Cho, A., et al. (2019). Hierarchical organization of cortical and thalamic connectivity. Nature 575, 195–202.

Hubel, D.H., and Wiesel, T.N. (1967). Cortical and callosal connections concerned with the vertical meridian of visual fields in the cat. J. Neurophysiol. 30, 1561–1573.

Imig, T.J., and Brugge, J.F. (1978). Sources and terminations of callosal axons related to binaural and frequency maps in primary auditory cortex of the cat. J. Comp. Neurol. 182, 637–660.

Isaacson, J.S., and Scanziani, M. (2011). How inhibition shapes cortical activity. Neuron 72, 231–243.

Karayannis, T., Huerta-Ocampo, I., and Capogna, M. (2007). GABAergic and pyramidal neurons of deep cortical layers directly receive and differently integrate callosal input. Cereb. Cortex 17, 1213–1226.

Kato, H.K., Gillet, S.N., and Isaacson, J.S. (2015). Flexible Sensory Representations in Auditory Cortex Driven by Behavioral Relevance. Neuron 88, 1027–1039.

Kitzes, L.M., and Doherty, D. (1994). Influence of callosal activity on units in the auditory cortex of ferret (Mustela putorius). J. Neurophysiol. 71, 1740–1751.

Lee, C.C., and Winer, J.A. (2008). Connections of cat auditory cortex: II. Commissural system. J. Comp. Neurol. 507, 1901–1919.

Lee, A.T., Gee, S.M., Vogt, D., Patel, T., Rubenstein, J.L., and Sohal, V.S. (2014). Pyramidal neurons in prefrontal cortex receive subtype-specific forms of excitation and inhibition. Neuron 81, 61–68.

Lee, K.-S., Vandemark, K., Mezey, D., Shultz, N., and Fitzpatrick, D. (2019). Functional Synaptic Architecture of Callosal Inputs in Mouse Primary Visual Cortex. Neuron 101, 421–428.e5.

Li, L., Li, Y., Zhou, M., Tao, H.W., and Zhang, L.I. (2013). Intracortical multiplication of thalamocortical signals in mouse auditory cortex. Nat. Neurosci. 16, 1179–1181.

Lien, A.D., and Scanziani, M. (2013). Tuned thalamic excitation is amplified by visual cortical circuits. Nat. Neurosci. 16, 1315–1323.

Manzoni, T., Barbaresi, P., Conti, F., and Fabri, M. (1989). The callosal connections of the primary somatosensory cortex and the neural bases of midline fusion. Exp.Erimental Brain Res. 76, 251–266.

Mitani, A., and Shimokouchi, M. (1985). Neuronal connections in the primary auditory cortex: An electrophysiological study in the cat. J. Comp. Neurol. 235, 417–429.

Pachitariu, M., Steinmetz, N.A., Kadir, S.N., Carandini, M., and Harris, K.D. (2016). Fast and accurate spike sorting of high-channel count probes with KiloSort. Adv. Neural Inf. Process. Syst. 29, 4448–4456.

Payne, B., Elberger, A., Berman, N., and Murphy, E. (1980). Binocularity in the cat visual cortex is reduced by sectioning the corpus callosum. Science (80-.). 207, 1097–1099.

Petreanu, L., Huber, D., Sobczyk, A., and Svoboda, K. (2007). Channelrhodopsin-2-assisted circuit mapping of long-range callosal projections. Nat Neurosci 10, 663–668.

Pettersen, K.H., Devor, A., Ulbert, I., Dale, A.M., and Einevoll, G.T. (2006). Current-source density estimation based on inversion of electrostatic forward solution: Effects of finite extent of neuronal activity and conductivity discontinuities. J. Neurosci. Methods 154, 116–133.

Rock, C., and Apicella, A. j. (2015). Callosal Projections Drive Neuronal-Specific Responses in the Mouse Auditory Cortex. J. Neurosci. 35, 6703–6713.

Rock, C., Zurita, H., Lebby, S., Wilson, C.J., and Apicella, A. junior (2018). Cortical Circuits of Callosal GABAergic Neurons. Cereb. Cortex 28, 1154–1167.

Rossant, C., Kadir, S.N., Goodman, D.F.M., Schulman, J., Hunter, M.L.D., Saleem, A.B., Grosmark, A., Belluscio, M., Denfield, G.H., Ecker, A.S., et al. (2016). Spike sorting for large, dense electrode arrays. Nat. Neurosci. 19, 634–641.

Rouiller, E.M., Simm, G.M., Villa, A.E.P., de Ribaupierre, Y., and de Ribaupierre, F. (1991). Auditory corticocortical interconnections in the cat: evidence for parallel and hierarchical arrangement of the auditory cortical areas. Exp. Brain Res. 86, 483–505.

Schmidt, K.E., Lomber, S.G., and Innocenti, G.M. (2010). Specificity of Neuronal Responses in Primary Visual Cortex Is Modulated by Interhemispheric CorticoCortical Input. Cereb. Cortex 20, 2776–2786.

Siegle, J.H., López, A.C., Patel, Y.A., Abramov, K., Ohayon, S., and Voigts, J. (2017). Open Ephys: An open-source, plugin-based platform for multichannel electrophysiology. J. Neural Eng. 14.

Silver, R.A. (2010). Neuronal arithmetic. Nat. Rev. Neurosci. 11, 474–489.

Sturgill, J.F., and Isaacson, J.S. (2015). Somatostatin cells regulate sensory response fidelity via subtractive inhibition in olfactory cortex. Nat. Neurosci. 18, 531–535.

Tolhurst, D.J., Movshon, J.A., and Dean, A.F. (1983). The statistical reliability of signals in single neurons in cat and monkey visual cortex. Vision Res. 23, 775–785.

Wise, S.P., and Jones, E.G. (1976). The organization and postnatal development of the commissural projection of the rat somatic sensory cortex. J. Comp. Neurol. 168, 313–343.

Wunderle, T., Eriksson, D., Peiker, C., and Schmidt, K.E. (2015). Input and output gain modulation by the lateral interhemispheric network in early visual cortex. J. Neurosci. 35, 7682–7694.

